# Lactate Promotes an Anti-Inflammatory Phenotype in Activated Microglia

**DOI:** 10.64898/2026.09.25.754340

**Authors:** Ifrah Omar Ibrahim, Anne-Karine Bouzier-Sore, Jane Frezals, Pierre Goudeneche, Hélène Roumes, Jeanny Laroche-Traineau

**Author notes:** Corresponding author: Anne-Karine Bouzier-Sore, Univ. Bordeaux, CNRS, CRMSB, UMR 5536, F-33000 Bordeaux, France. These authors contributed equally to this work (Ifrah Omar Ibrahim, Anne-Karine Bouzier-Sore). These authors are sharing the last position (Hélène Roumes, Jeanny Laroche-Traineau).

## Abstract

Microglial activation is a central component of neuroinflammatory responses in many brain pathologies. Increasing evidence indicates that microglial phenotype is tightly linked to cellular metabolism, with pro-inflammatory activation associated with enhanced glycolytic flux. Lactate, traditionally considered a metabolic substrate, has recently emerged as a signaling molecule capable of modulating immune responses. However, its direct impact on microglial inflammatory activation remains incompletely understood.

In the present study, we investigated the effects of lactate on microglial phenotype under inflammatory conditions using primary rat microglial cultures stimulated with lipopolysaccharide (LPS). Microglial activation was assessed through the expression of phenotypic markers, cytokine production, and secreted chemokine profiles. LPS stimulation induced a strong pro-inflammatory response characterized by increased CD86 expression, elevated TNF-α secretion, and enhanced release of several pro-inflammatory chemokines. Post-treatment with sodium L-lactate significantly attenuated these inflammatory responses, reducing pro-inflammatory marker expression and cytokine secretion, while restoring the anti-inflammatory marker CD206.

To explore the relevance of these findings in a pathological context, the effects of lactate were further examined in a neonatal rat model of hypoxia-ischemia. Sodium L-lactate administration after injury reduced microglial activation and promoted a shift toward an anti-inflammatory phenotype in cortical regions, whereas hippocampal microglia showed a more limited response.

Together, these results demonstrate that lactate directly modulates microglial inflammatory activation and cytokine production *in vitro* and suggest that lactate-mediated metabolic signaling may contribute *in vivo* to the regulation of neuroinflammatory responses.

**Graphical abstract:** 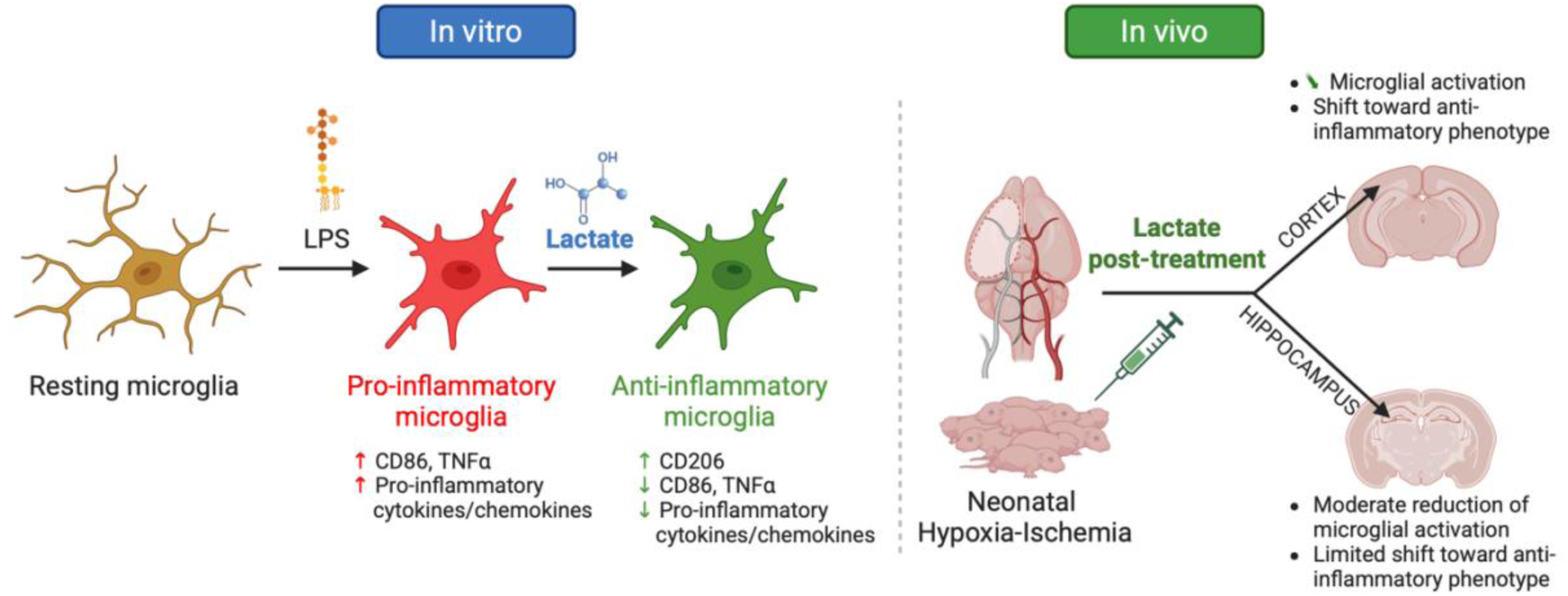

**Highlights:**

- Lactate reduces pro-inflammatory activation in primary microglia.
- Lactate promotes anti-inflammatory microglial phenotype *in vitro*.
- Cortical microglia respond better to lactate than hippocampal microglia.

## Introduction

Neonatal hypoxia-ischemia (NHI) is one of the major causes of perinatal death and acquired neurobehavioral and cognitive sequelae. Its incidence rate of 1-8‰ in developed countries makes it a major public health issue. NHI results from a reduction in cerebral blood flow leading to restricted oxygen and energy substrate supply to the brain. Brain injury arises from complex processes involving multiple mechanisms and signaling pathways. One of the main pathogenic factors is inflammation, induced by the activation of the central immune system, microglia, and peripheral immune cells (macrophages, monocytes and mast cells) (Ziemka-Nalecz M, *et al*., 2017). At the central nervous system level, reactive astrocytes may contribute to inflammation (Liddelow SA and Barres BA, 2017; Sen E and Levison SW, 2006; Sullivan SM, *et al*., 2010). However, the role of astrocytes in the progression of ischemic brain injury, particularly in neonates, remains controversial. As highlighted by Liddelow and Sofroniew (2019), reactive astrocytes can exert both detrimental and protective effects by releasing pro- and anti-inflammatory mediators, thereby influencing the survival of surrounding neurons (Jiao M, *et al*., 2020; Jin D, *et al*., 2024; Liddelow SA, *et al*., 2024; Liddelow SA and Sofroniew MV, 2019). Studies suggest that the deleterious inflammatory response observed in NHI is thought to be mainly due to microglial activation (Li B, *et al*., 2017; Serdar M, *et al*., 2019). Inflammatory signals are generated within minutes after NHI and can extend for several weeks (Fleiss B, *et al*., 2021). Furthermore, microglia are among the first cells to become activated after NHI (Millar LJ, *et al*., 2017). Under physiological conditions, microglial cells display a characteristic ramified morphology, acting as sensors to dynamically monitor the microenvironment and maintain brain homeostasis (Nimmerjahn A, *et al*., 2005). In the context of cerebral injury, such as NHI, microglia transform into reactive cells, adopting an amoeboid shape (Green TRF and Rowe RK, 2024). These activated microglia migrate to the injury site, proliferate, and shift toward a pro-inflammatory phenotype, triggering the release of cytokines, chemokines, reactive oxygen species (ROS), and matrix metalloproteinases that contribute to neuronal death. This neuronal death impairs brain development and underlies long-term cognitive and behavioral disorders (Dommergues MA, *et al*., 2003; Hagberg H, *et al*., 2012). It has been shown that reducing microglial activation after NHI is associated with decreased brain damage (Dommergues MA, *et al*., 2003); however, complete inhibition of this activity, for instance by intracerebral clodronate injection in 7-day-old pups (P7), or the use of transgenic mice, can exacerbate lesion severity in neonatal stroke (Faustino JV, *et al*., 2011; Lalancette-Hebert M, *et al*., 2007). These findings indicate that microglial activation is not uniformly detrimental and that both pro- and anti-inflammatory responses may contribute to the outcome following NHI. Accordingly, the therapeutic challenge is not to suppress microglial activation altogether, but rather to limit excessive pro-inflammatory responses while preserving beneficial functions involved in synaptic pruning, tissue repair, and recovery. These context-dependent functions are supported by the phenotypic plasticity of microglia, which can adopt distinct functional states in response to the surrounding microenvironment.

Microglia with an anti-inflammatory phenotype exhibit immunosuppressive and neuroprotective properties through the release of anti-inflammatory and neurotrophic factors that promote tissue remodeling and neuronal repair (Ekdahl CT, *et al*., 2009), whereas activated microglia can also contribute to neuroprotection by eliminating apoptotic debris and supporting tissue restoration and resolution of inflammation. Importantly, these phenotypic states are not fixed but are closely linked to metabolic reprogramming. In their homeostatic state, microglia predominantly rely on oxidative metabolism to meet their energy demands, whereas pro-inflammatory activation is associated with a metabolic switch towards glycolysis. This metabolic plasticity enables microglia to adapt their energy production to changes in nutrient availability and functional demands. In particular, pro-inflammatory microglia increase their reliance on glycolysis, as reflected by enhanced hexokinase and lactate dehydrogenase activity and increased expression of the glucose transporter GLUT1 (Gimeno-Bayon J, *et al*., 2014). In contrast, anti-inflammatory microglia predominantly generate ATP through oxidative phosphorylation (Bernier LP, *et al*., 2020; Voloboueva LA, *et al*., 2013). Modulating this metabolic switch may thus influence the microglial phenotype, reducing neuronal death and preserving cognitive functions.

A better understanding of microglial activation mechanisms is essential to clarify the deleterious effects of NHI and to optimize neuroprotective strategies aimed at preserving brain function. Our previous *in vivo* studies have demonstrated the neuroprotective effect of sodium L-lactate administration after NHI (Roumes H, *et al*., 2021; Roumes H, *et al*., 2020). Building on these *in vivo* results, the present work aimed to decipher, at the cellular level, the effects of sodium L-lactate on microglial phenotype under inflammatory conditions. We first evaluated the impact of sodium L-lactate on primary microglial cultures stimulated with lipopolysaccharide (LPS), by assessing CD206, CD86, and TNF-α expression. CD206 (Mannose Receptor) is commonly associated with homeostatic/anti-inflammatory microglia states, whereas CD86 is associated with pro-inflammatory microglia states, and TNF-α is a hallmark pro-inflammatory cytokine (Jurga AM, *et al*., 2020). We then assessed the effects of sodium L-lactate on *ex vivo* brain slices from neonatal rats subjected to NHI, by analyzing CD206 and CD86 expression levels, as well as microglia morphology. Our findings indicate that post-insult sodium L-lactate administration modulates microglial responses, promoting a shift toward an anti-inflammatory phenotype while limiting pro-inflammatory activation, with a stronger effect in cortical regions than in the hippocampus, suggesting the regional differences may influence microglial responsiveness.

## Experimental procedures

### Primary microglia Culture

Primary microglia were isolated from the cortices of neonatal Wistar rats (P1-P3). Following decapitation, performed in accordance with the European Community Council Directive of November 24, 1986 (86/609/EEC), meninges were carefully removed and both hemispheres dissected in ice-cold PBS (Phosphate-Buffered Saline) 1X, glucose (1 g/l) (Merck, St Quentin Fallavier, France), 100 units/mL penicillin, 100 μg/mL streptomycin, and 250 ng/mL Amphotericin B (antibiotic-antimycotic, Gibco 15240-096, Fisher Scientific). Cortical tissue was mechanically dissociated using Dumont forceps and digested for 15 min at 37 °C in a solution containing trypsin (0.25%, Gibco) and DNase I (60 µg/mL, Sigma-Aldrich). The digestion was stopped by adding Dulbecco’s Modified Eagle Medium (DMEM; Gibco 15313165, Fisher Scientific) supplemented with 10% fetal bovine serum (FBS), penicillin (100 U/mL), and streptomycin (100 µg/mL) (Gibco 15240-062). The suspension was triturated 15 times with a 5 mL pipette, then 20 times with a 2 mL pipette, and centrifuged at 500 g for 5 min. The pellet was further mechanically dissociated by triturating 20 times with a 1 mL pipette and 20 times with a 200 µL pipette. Cells were resuspended in 10 mL DMEM supplemented with 10% FBS, penicillin (100 U/mL), and streptomycin (100 μg/mL), and plated at a density of 5 × 10⁵ cells per well in 100 mm culture dishes (Fisher Scientific) pre-coated with poly-L-lysine (100 µg/mL; Sigma-Aldrich). Cultures were maintained at 37 °C in a humidified 5% CO₂ atmosphere, and the medium was replaced every 4 days. After 12 days in vitro, microglia were detached by orbital shaking at 37 °C for 1 h at 160 rpm, following protocols adapted from Lin *et al*. (Lin L, *et al*., 2017) and Miličević *et al*. (Milicevic K, *et al*., 2022). Floating microglia were gently collected, counted, and re-plated for subsequent experiments (see below).

### Cell Treatments

Primary microglia were seeded at a density of 1 × 10⁵ cells per well (1 mL) in 24-well plates (Corning® Primaria™, Dutcher, Bernolsheim, France) with DMEM, low glucose 1g/L (Gibco 15313165, Fisher Scientific), 10% heat-inactivated FBS, penicillin, and streptomycin (Gibco, 15240-096, Fisher Scientific). Cultures were maintained at 37 °C in a humidified atmosphere with 5% CO₂. Cells were stimulated with 50 ng/mL LPS (Escherichia coli serotype 0127:B8, Merck) to mimic the inflammatory conditions of neonatal hypoxic-ischemic injury. The effects of sodium L-lactate (Merck, L7022-5G) at 5 mM were assessed. Table 1 presents the different experimental conditions over 48 h. Experiments were performed in duplicate for each condition, and the number of independent biological replicates (n) is indicated in the statistical plots. Technical replicates were averaged prior to statistical analysis.

**Table 1:**
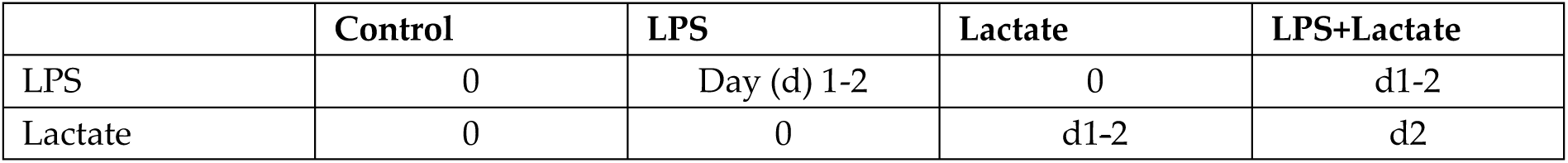
Cell treatment over 48 h.

### Primary microglia immunofluorescence

Primary microglia were seeded at 1 × 10⁵ cells/well on sterilized 12 mm glass coverslips in 24-well plates (Corning®) and cultured in DMEM supplemented with 10% FBS, 100 U/mL penicillin, and 100 μg/mL streptomycin. Conditioned media were collected, centrifuged at 400 × g for 10 min, and stored at −20 °C for subsequent TNF-α quantification by ELISA. Cells were rinsed with 1 mL of 1X PBS and fixed with 4% paraformaldehyde (PFA) (15670799, Fisher Scientific) for 20 min at room temperature, then rinsed again with 1X PBS. After permeabilization of cell membranes in PBS buffer containing 0.25% (v/v) Triton X-100 for 15 min, blocking was performed using 2% (w/v) BSA buffer with 0.25% (v/v) Triton X-100 for 1 h. Cells were incubated overnight at 4 °C with primary antibodies: anti-Iba1 (Abcam, ab48004) to confirm microglial identity, and anti-GFAP (Proteintech, 16825-1-AP) to exclude astrocytic contamination, and anti-CD86 (Proteintech, 13395-1-AP) or anti-CD206 (Abcam, ab64693) to identify pro-inflammatory and anti-inflammatory phenotypes, respectively. Following three washes with 1X PBS containing 0.05% Triton X-100, appropriate secondary antibodies (Alexa Fluor 488 or 568, Abcam) were applied for 1 h. Coverslips were mounted with Vectashield containing DAPI (4′, 6-diamidino-2-phenylindole, Vector Laboratories) and imaged using a Nikon Eclipse 90i microscope (20× and 40×). Fluorescence intensity was quantified using Fiji/ImageJ software.

### Secreted cytokine profiling by proteome profiler antibody array

To assess the secreted cytokine profiles under different conditions, culture supernatants from primary microglia were collected and centrifuged at 2,500 × g for 5 min. Supernatants were analyzed using the Rat Cytokine Array Panel A (Proteome Profiler Antibody Array, ARY008, Bio-Techne) following the manufacturer’s instructions. Briefly, diluted culture supernatants were mixed with a cocktail of biotinylated detection antibodies and incubated overnight at 4°C on nitrocellulose membranes pre-spotted with capture antibodies specific for target cytokines. After three washes, IRDye 800CW Streptavidin (Licor Biosciences 926-32230, Fisher Scientific) diluted 1:2,000 was added and incubated for 30 min at room temperature. Signals were detected using the Odyssey Imager at 800 nm.

### TNF-α quantification by ELISA

Cell culture supernatants were collected and centrifuged at 400 × g for 10 min at 20°C. TNF-α levels were measured using a Rat TNF-α Mini ABTS ELISA Development Kit (900-M73, PeproTech, Neuilly-sur-Seine, France) according to the manufacturer’s instructions. Briefly, 100 μL of capture antibody (2 μg/mL) was added to each well of a 96-well plate (Costar™ 3590, Dutcher) and incubated overnight at 4°C. Plates were washed twice with PBS-BSA (0.1% w/v) containing 0.05% Tween-20, and nonspecific binding sites were blocked with PBS-BSA (1% w/v) for 1 h at room temperature. After washing, a standard curve of TNF-α (31.25–4,000 pg/mL) was applied in duplicate, and diluted culture supernatants (1:2 in PBS-BSA 0.1% w/v, 0.05% Tween-20) were added in duplicate. Plates were incubated for 2 h, washed four times, and biotinylated detection antibody (0.5 μg/mL) was added for 2 h. Following four washes, Avidin-horseradish peroxidase (HRP) conjugate (1:2,000) was applied for 30 min. After three washes, 100 μL of substrate solution (0.4 mg/mL o-phenylenediamine dihydrochloride (OPD), 0.4 mg/mL urea hydrogen peroxide in 0.05 M citrate-phosphate buffer, pH 5.0) was added and incubated for 15 min in the dark. The reaction was stopped with 50 μL of 2 N sulfuric acid, and absorbance was measured at 490 nm using a microplate reader (SpectraMax, Molecular Devices).

### Animals

All animal procedures were performed in compliance with the European Communities Council Directive of 24 November 1986 (86/609/EEC) on animal experimentation. Experimental protocols adhered to the ethical guidelines of the French Ministry of Agriculture and Forestry as well as the ARRIVE guidelines, and were approved by the Bordeaux Ethical Committee for Animal Research (C2EA-50, authorization n° 37955). Pregnant Wistar RJ-HAN females (Janvier Laboratories, France; n = 3) were received at gestational day 16 and housed under a 12:12-hour light/dark cycle with free access to food and water.

### Induction of NHI and experimental groups

Neonatal Wistar rats at P7, of both sexes, were used to induce hypoxic-ischemic injury. Animals were anesthetized with isoflurane (4% induction, 1.5% maintenance), and a midline neck incision was performed under local anesthesia (0.5% lidocaine). The left common carotid artery was carefully exposed and permanently ligated using a 7–0 silk suture. Surgical procedures were kept brief (6–7 min, not exceeding 14 min) to minimize risk of cardiac or respiratory complications. Following surgery, pups were allowed to recover for 30 min on a heated mattress to maintain normothermia. For hypoxic exposure, pups were placed in a humidified chamber containing 8% oxygen balanced with 92% nitrogen for 2 h. The chamber was submerged in a 36 ± 1 °C water bath (Intensive Care Unit Warmer, Harvard Apparatus) to ensure stable body temperature. Sham-operated pups were separated from the dam and maintained in a heated environment (33 ± 1 °C) for the same duration. After hypoxic exposure, animals were returned to the heated mattress before being reunited with the dam. Rectal temperatures confirmed maintenance of normothermia throughout the procedure. Three experimental groups were established, with pups originating from three different dams randomly assigned to one of the groups: (i) Sham group, in which P7 pups received an intraperitoneal injection of 0.9% NaCl 150 min after sham surgery and once daily for the following 48 h (n = 6); (ii) HIC group (Hypoxia-Ischemia Control), in which P7 pups underwent the NHI procedure and received intraperitoneal injections of 0.9% NaCl at 150 min post-ligation and once daily for the next 48 h (n = 6); and (iii) HIL group (Hypoxia-Ischemia + Lactate), in which P7 pups underwent the NHI procedure and received intraperitoneal injections of sodium L-lactate (1 mol/L, 4 μl/g body weight) at 150 min, 24 h, and 48 h after carotid ligation (n = 6).

### Brain Collection for Histology and Immunohistochemistry

At P9, pups (n = 6 per experimental group) were deeply anesthetized with a ketamine-xylazine mixture. Animals underwent intracardiac perfusion first with PBS (50 mL over 10 min) to clear the vasculature, followed by 50 mL of 4% PFA for fixation. Brains were carefully removed and post-fixed overnight in 4% PFA at 4 °C. Subsequently, tissues were cryoprotected in 30% sucrose in PBS for 48 h at 4 °C. After cryoprotection, brains were frozen using nitrogen vapor and stored at −80 °C until sectioning with a cryostat. To assess cellular microglial activation, and polarization, 16-μm cryostat sections from the hippocampus and cortex were prepared. Sections were first thawed at 37 °C for 15 min and washed six times in PBS over a 30-min period. They were then blocked in PBS containing 3% (w/v) BSA and 0.3% (v/v) Triton X-100 (Merck) for 1.5 h at room temperature. Primary antibodies (see Table 1) were applied in blocking buffer and incubated overnight at 4 °C. After rinsing six times in PBS for 30 min, sections were incubated for 1 h at room temperature with appropriate Alexa Fluor-conjugated secondary antibodies: goat-targeting (1:500, Alexa Fluor 488, #Ab150129) and rabbit-targeting (1:500, Alexa Fluor 568, #Ab175470). Following six additional PBS washes, slides were mounted with VECTASHIELD containing DAPI (Vector Laboratories, Eurobio, France) for nuclear counterstaining. Fluorescence images were acquired using 20× and 40× objectives on a Nikon Eclipse 90i microscope.

**Table 2:** Primary antibodies.

| Host/ Primary antibody | Producer-Reference | Anticorps Type | Dilution |
| --- | --- | --- | --- |
| Goat/ Anti-Iba1 | Abcam ab48004 | Monoclonal | 1:1000 |
| Rabbit/Anti-CD86 | Proteintech 13395-1-AP | Polyclonal | 1:4000 |
| Rabbit/ Anti-Mannose receptor (CD206) | Abcam ab64693 | Polyclonal | 1:10000 |

### Microglial Morphology Analysis

Microglial morphology was assessed using Iba1 immunofluorescence images. Images were first opened and converted to 8-bit format in Fiji/ImageJ software. A threshold was applied to distinguish microglial structures from the background, and images were subsequently binarized and skeletonized using the “Binary Skeleton” tool. To remove small artifacts, the “Analyze Particles” function was applied with a size range of 50 pixels to infinity. Skeletonized images were then analyzed using the Skeleton Analysis plugin to quantify morphological parameters, including the number of branches, junctions, and triple points. All measurements were performed under consistent image processing settings to ensure comparability between conditions.

### Statistical analysis

All measurements were performed blindly by two independent experimenters, with intra-experimenter variability below 1% and inter-experimenter variability below 5%. Statistical analyses and graph generation were performed using GraphPad Prism software (version 7.00, GraphPad Software). Data are presented as mean ± standard error of the mean (SEM), and the number of cultures or animals included in each experiment is indicated in the figures using dot plots. Dataset distributions were evaluated with the Shapiro–Wilk normality test. For normally distributed data, intergroup differences were assessed using one-way analysis of variance (ANOVA) followed by Holm-Sidak multiple comparisons correction. Non-normally distributed data were analyzed using the Kruskal-Wallis test with Dunn’s post hoc multiple comparisons correction. Statistical significance was defined as p < 0.05, and the following notation is used throughout: *p < 0.05, **p < 0.01, ***p < 0.001, and ****p < 0.0001.

## Results

### Post-LPS sodium L-lactate treatment promotes an anti-inflammatory phenotype in primary microglia

Primary microglial cultures displayed a high level of purity (92%), ensuring that the measured responses predominantly reflected microglial activation (data not shown).

As expected, LPS stimulation induced a robust pro-inflammatory shift. CD86 expression markedly increased (+144 ± 17% *vs*. control; Figure 1A-B), confirming the acquisition of a pro-inflammatory phenotype. Post-LPS sodium L-lactate treatment significantly attenuated this response, reducing CD86 expression to near-baseline levels (+23 ± 29% *vs*. control), indicating a partial reversal of microglial activation.

**Figure 1:**
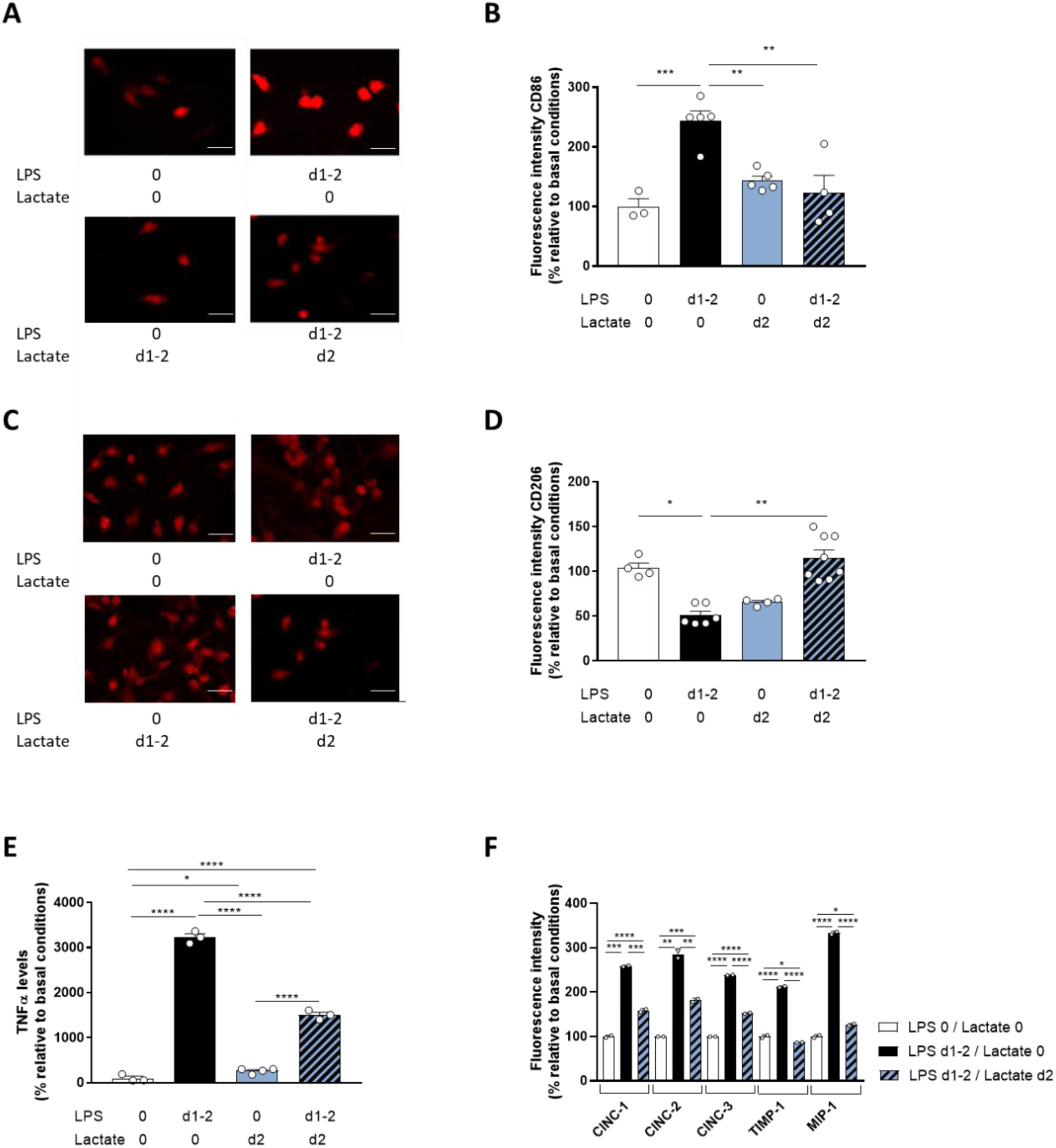
Effect of 24h-lactate treatment on primary microglia culture on 48h-LPS stimulation. (A) Representative images of CD86 immunofluorescence in the 4 experimental conditions: control, 48h-LPS treatment (on days 1 and 2 – d1-2), 24h-lactate treatment (on day 2 – d2), and the combination 48h-LPS (d1-2) + 24h-lactate (d2). (B) Quantification of CD86 fluorescence intensity, expressed as a % relative to the control. (C) Representative image of CD206 immunofluorescence in the 4 conditions. (D) Quantification of CD206 fluorescence intensity, expressed as a % relative to the control. (E) TNF-α levels measured by ELISA, in the 4 experimental conditions, expressed as a % relative to control. (F) Effect of lactate on immunoreactivity of the secreted cytokine profiles following LPS stimulation. Scale bar: 50 µm. Data are presented as mean ± SEM. Each dot represents one independent culture. *p < 0.05, **p < 0.01, ***p < 0.001, ****p < 0.0001 (Kruskal-Wallis (B), ANOVA (D, E, F), with Dunn’s or Holm-Sidak comparison correction.

Conversely, the anti-inflammatory marker CD206 was significantly downregulated by LPS (−49 ± 5% *vs*. control; Figure 1C-D). Sodium L-lactate post-treatment effectively restored CD206 expression, supporting a shift toward an anti-inflammatory phenotype.

TNF-α production exhibited a strong induction following LPS exposure (+3130 ± 78% *vs*. control; Figure 1E). Sodium L-lactate administration markedly reduced TNF-α levels, further confirming its ability to mitigate the LPS-induced inflammatory response.

To complement these marker-based analyses, a Proteome Profiler Antibody Array was performed to assess the profile of secreted chemoattractant cytokines and chemokines (Figure 1F). LPS stimulation increased the secretion of several pro-inflammatory mediators, including CINC-1 (Cytokine-Induced Neutrophil Chemoattractant), CINC-2, CINC-3, TIMP 1 (Tissue Inhibitor of Metalloproteinases-1), and MIP-1α (Macrophage Inflammatory Protein-1α). Notably, sodium L-lactate post-treatment markedly attenuated the production of all these cytokines, confirming that lactate not only modulates intracellular inflammatory markers but also effectively suppresses the secretion of key chemokines and cytokines involved in microglial activation and leukocyte recruitment.

### Sodium L-lactate modulates microglial activation in vivo following hypoxic-ischemic event

Microglial activation was first evaluated 48 h after NHI using Iba1 immunoreactivity (Figure 2A-B). In the cortex, Iba1 levels were increased in the control group (HIC) compared to Sham animals (mean: 726 ± 64% relative to the Sham group). Post-NHI administration of sodium L-lactate (HIL group) normalized cortical Iba1 expression, restoring it to values comparable to Sham (mean for HIL group: 242 ± 50%). In the CA1 hippocampal region, NHI similarly induced a significant increase in Iba1 immunoreactivity (mean for HIC group: 748 ± 29% relative to the Sham group; Figure 2C-D). Sodium L-lactate treatment significantly reduced this increase (mean for HIL group: 312 ± 9% relative to the Sham group), although Iba1 levels remained higher than in Sham animals. Consistent with these changes in Iba1 expression, NHI also induced a marked increase in microglial cell numbers both in the cortex (353 ± 5% relative to the Sham group; Figure 2E-G) and in the hippocampus (687 ± 25% relative to the Sham group; Figure 2F-H). Sodium L-lactate treatment significantly attenuated microglial recruitment in the cortex (Figure 2E) and partially reduced it in the CA1 region, although this reduction did not reach statistical significance compared with the HIC group (Figure 2F).

**Figure 2:**
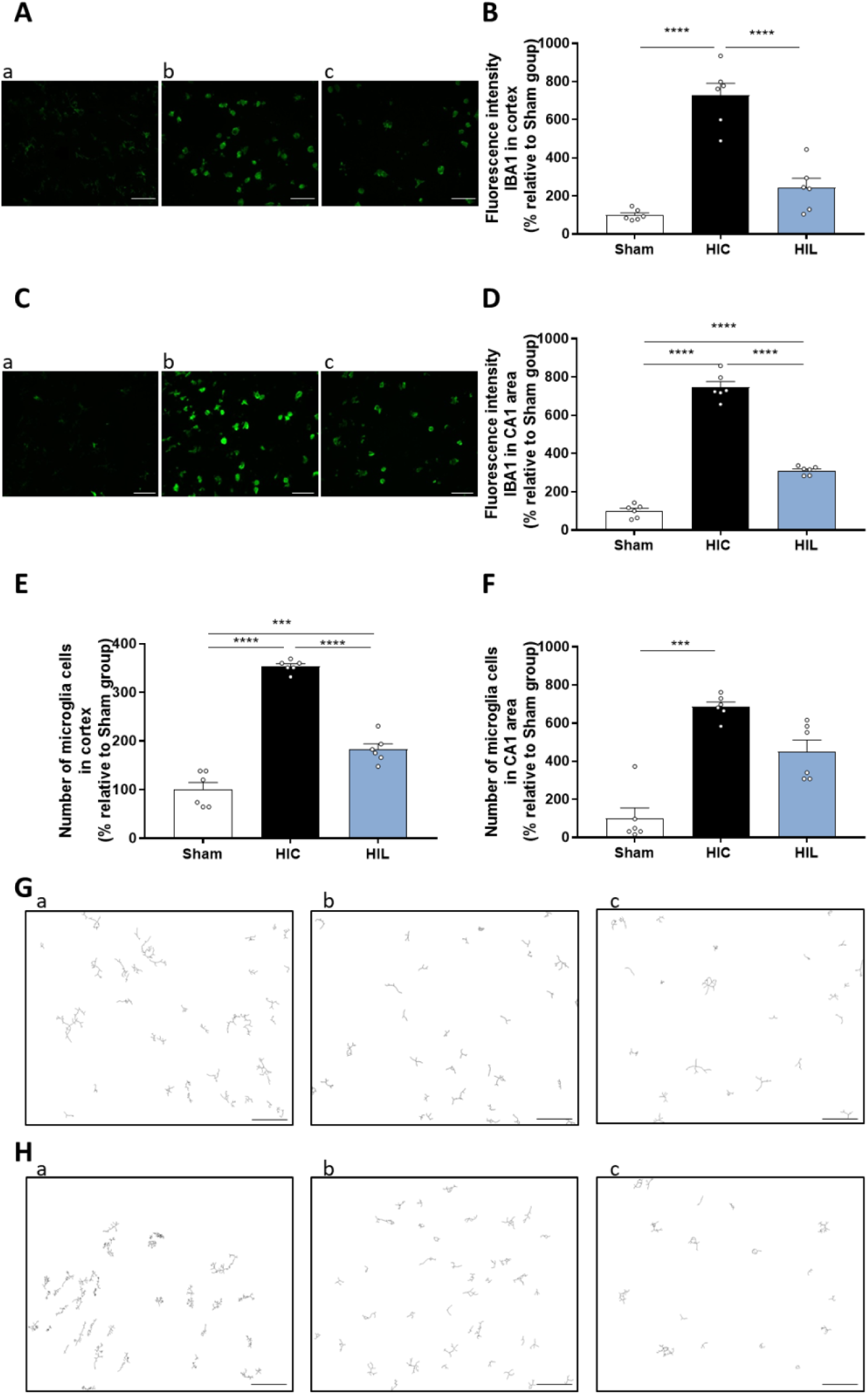
Effect of three consecutive daily i.p. administrations (d0-d1-d2) of Na-L-lactate following NHI on microglia activation at d2. (A) Representative images of Iba1 immunofluorescence in the cortex for the 3 groups 48h after the insult: (a) Sham, (b) HIC, and (c) HIL. (B) Quantification of Iba1 fluorescence intensity in the cortex, expressed as a % relative to the Sham group. (C) Representative images of Iba1 immunofluorescence in the CA1 area for the 3 groups: (a) Sham, (b) HIC, and (c) HIL. (D) Quantification of Iba1 fluorescence intensity in the CA1 area, expressed as a % relative to the Sham group. (E) Number of Iba1-positive cells (microglia) in the cortex, expressed as a % relative to the Sham group. (F) Number of Iba1-positive cells in the CA1 area, expressed as a % relative to the Sham group. (G) Microglia skeletons in the cortex for (a) Sham, (b) HIC, and (c) HIL groups. (H) Microglia skeleton in the CA1 area for (a) Sham, (b) HIC, and (c) HIL groups. Images are representative of 6 independent pups/group. Scale bar: 50 µm. Data are presented as mean ± SEM. Each dot represents one independent animal. ***p < 0.001, ****p < 0.0001 (Kruskal-Wallis (F), ANOVA (B, D, E), with Dunn’s or Holm-Sidak comparison correction.

Microglial phenotype was characterized by assessing the expression of the pro-inflammatory marker CD86 and the anti-inflammatory/homeostatic marker CD206 (Figure 3). Pups underwent the hypoxia–ischemia procedure at P7 (defined as day 0 – d0), and brains were collected at d2 to assess microglial phenotype by immunofluorescence. Sodium L-lactate was administered by intraperitoneal injection at 3, 24, and 48 h after NHI.

**Figure 3:**
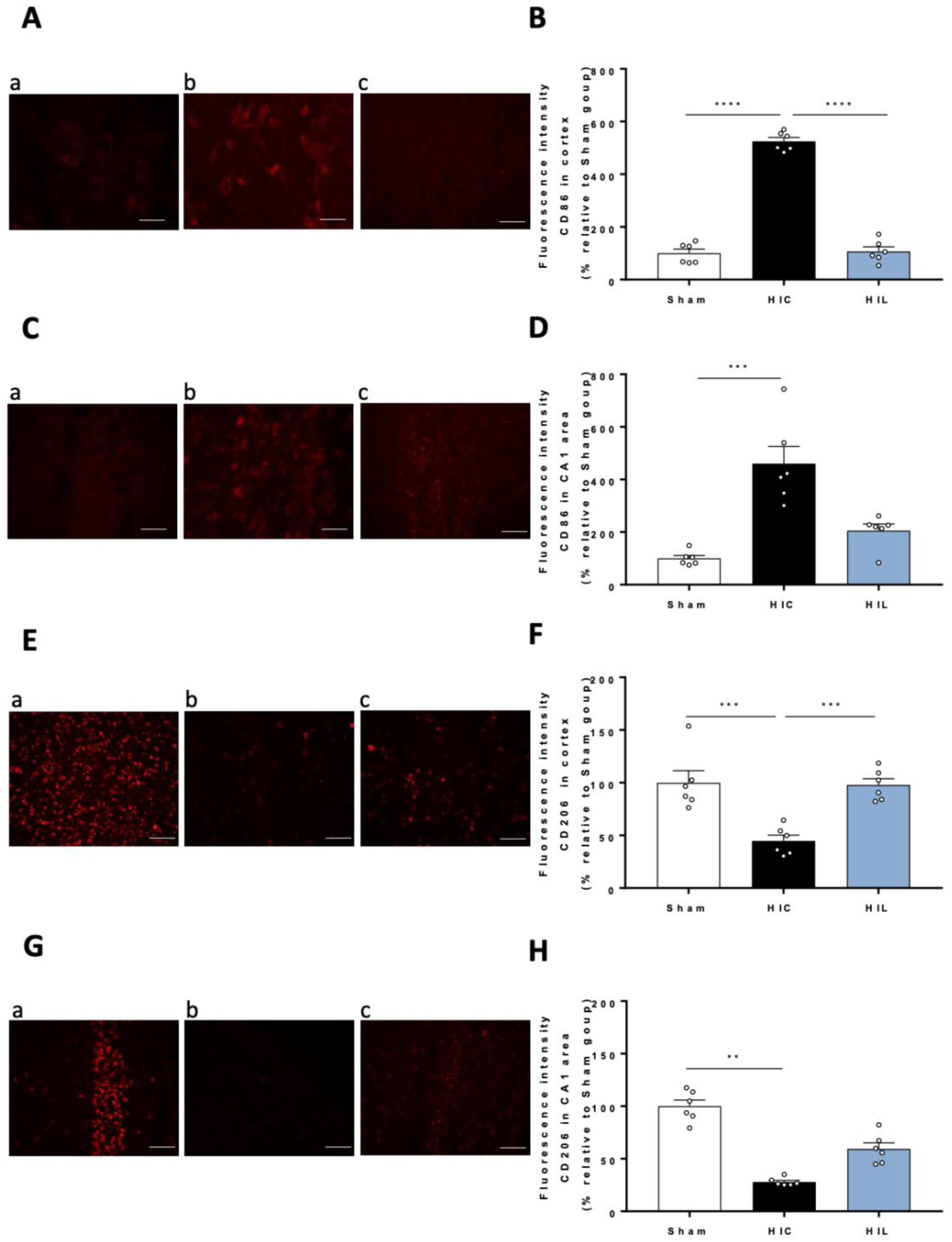
Effect of three consecutive daily i.p. administrations (d0-d1-d2) of Na-L-lactate following NHI on microglia inflammatory phenotype at d2. (A) Representative images of CD86 immunofluorescence in the cortex for the 3 groups: (a) Sham, (b) HIC, and (c) HIL. (B) Quantification of CD86 fluorescence intensity in the cortex, expressed as a % relative to the Sham group. (C) Representative images of CD86 immunofluorescence in the CA1 area for the 3 groups: (a) Sham, (b) HIC, and (c) HIL. (D) Quantification of CD86 fluorescence intensity in the CA1 area, expressed as a % relative to the Sham group. (E) Representative images of CD206 immunofluorescence in the cortex for the 3 groups: (a) Sham, (b) HIC, and (c) HIL. (F) Quantification of CD206 fluorescence intensity in the cortex, expressed as a % relative to the Sham group. (G) Representative images of CD206 immunofluorescence in the CA1 area for the 3 groups: (a) Sham, (b) HIC, and (c) HIL. (H) Quantification of CD206 fluorescence intensity in the CA1 area, expressed as a % relative to the Sham group. Images are representative of 6 independent pups/group. Scale bar: 100 µm. Data are presented as mean ± SEM. Each dot represents one independent animal. **p<0.01, ***p < 0.001, ****p < 0.0001 (Kruskal-Wallis (D, G), ANOVA (B, E), with Dunn’s or Holm-Sidak comparison correction.

Cortical CD86 expression was significantly upregulated following NHI (mean for HIC group: 525 ± 14% relative to the Sham group; Figure 3A-B), and this effect was fully inhibited by sodium L-lactate treatment, with CD86 expression returning to values comparable to those observed in Sham animals. In the hippocampus, however, NHI-induced CD86 upregulation was observed but was not significantly attenuated by sodium L-lactate treatment (460 ± 65% and 205 ± 25% relative to the Sham group, for HIC and HIL groups, respectively; Figure 3C-D).

Finally, CD206 expression was quantified as a marker associated with an anti-inflammatory and homeostatic microglial phenotype. Cortical CD206 levels were significantly reduced after NHI (45 ± 6% relative to the Sham group; Figure 3E-F), and sodium L-lactate treatment restored CD206 expression to values comparable to those observed in Sham animals. In contrast, hippocampal CD206 expression was also reduced following NHI (mean for HIC group: 28 ± 2% relative to the Sham group; Figure 3G-H), but the three consecutives daily sodium L-lactate administration failed to reverse this reduction, with CD206 expression reaching 59 ± 6% of Sham values in the HIL group.

To further characterize microglial activation, we assessed microglial morphological complexity using Skeleton Analysis (Figure 4). In the cortex, NHI induced a marked simplification of the microglial arborization, with a significant reduction in the number of branches compared with Sham animals (Figure 4A-B). This alteration was fully prevented by sodium L-lactate treatment. Similarly, the number of junctions, reflecting the complexity of the branching network, was significantly reduced following NHI and restored to Sham levels in the sodium L-lactate group (Figure 4C-D). Consistent with these findings, NHI also reduced the number of branching points (Figure 4E), whereas sodium L-lactate preserved microglial structural complexity.

**Figure 4:**
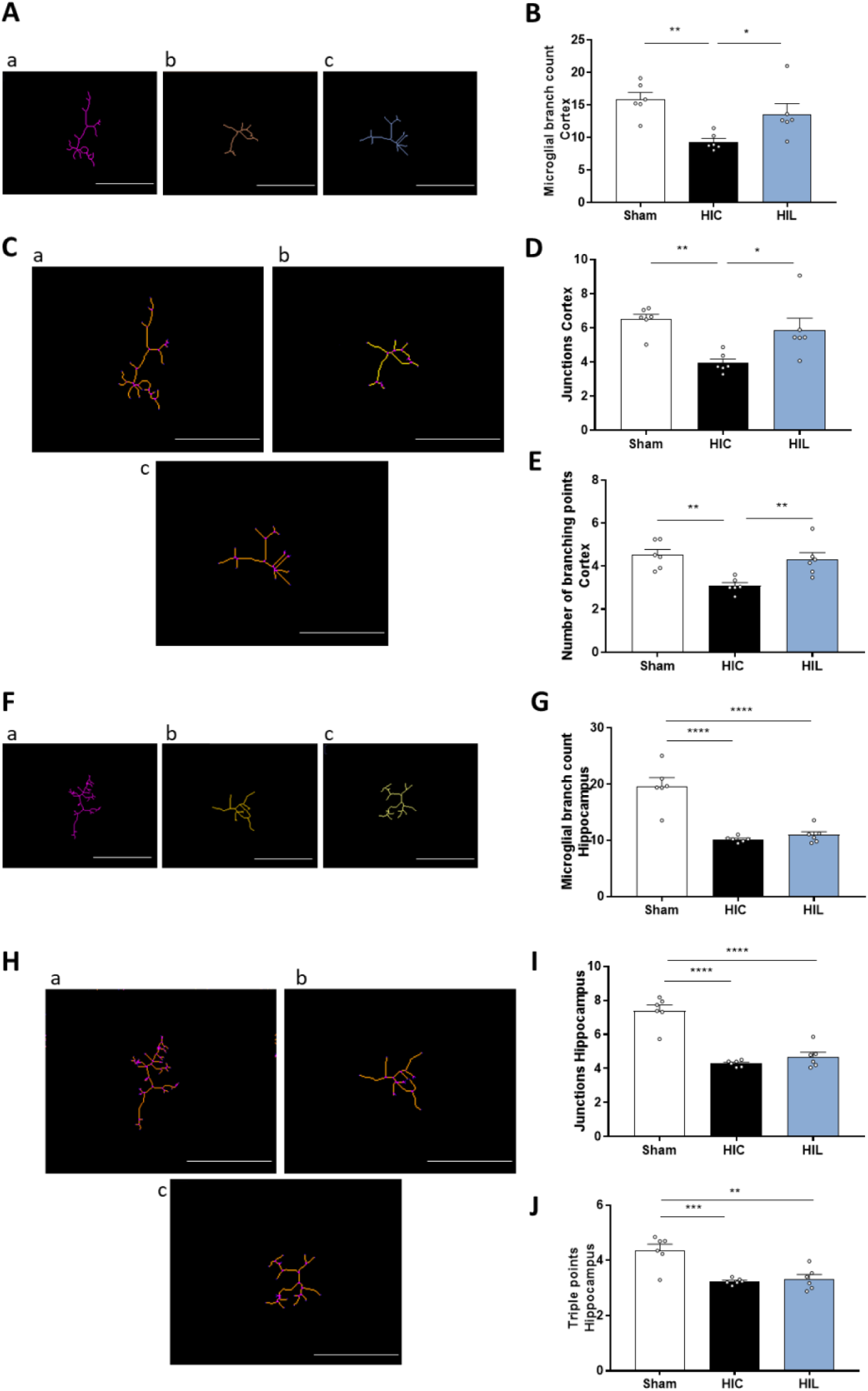
Effect of three consecutive daily i.p. administrations (d0-d1-d2) of Na-L-lactate following NHI on microglia inflammatory morphology at d2. (A) Representative images of microglia skeletons in the cortex, for the 3 experimental groups: (a) Sham, (b) HIC, and (c) HIL. (B) Microglial branch count in the cortex for the 3 experimental groups. (C) Representative images of skeletonized microglia, in the cortex, with junction’s points highlighted in violet, for the 3 groups: (a) Sham, (b) HIC, and (c) HIL. (D) Mean number of junctions in the cortex, for the 3 groups (E) Mean number of branching points in the cortex, for the 3 groups. (F) Representative images of microglia skeletons in the CA1 area, for the 3 groups: (a) Sham, (b) HIC, and (c) HIL. (G) Microglial branch count in the CA1 area for the 3 experimental groups. (H) Representative images of skeletonized microglia, in the CA1 area, with junction’s points highlighted in violet, for the 3 groups: (a) Sham, (b) HIC, and (c) HIL. (I) Mean number of junctions in the CA1 area, for the 3 groups. (J) Mean number of branching points in the CA1 area, for the 3 groups. Images are representative of 6 independent pups/group. Scale bar: 50 µm. Data are presented as mean ± SEM. Each dot represents one independent animal. *p < 0.05, **p <0.01, ***p < 0.001, ****p < 0.0001 ANOVA, with Holm-Sidak comparison correction.

In contrast, hippocampal microglia appeared more vulnerable to NHI-induced morphological alterations and less responsive to sodium L-lactate treatment. NHI significantly reduced the number of branches (Figure 4F-G), junctions (Figure 4H-I), and total branching points (Figure 4J). Sodium L-lactate treatment failed to restore any of these parameters, indicating a persistent reduction in microglial morphological complexity in the hippocampal CA1 region following NHI.

## Discussion

The present study demonstrated that lactate exerts an anti-inflammatory effect on microglia both *in vitro* and *in vivo* after NHI, supporting its potential role as a metabolic and immunomodulatory mediator.

In primary microglia, our findings suggest that post-LPS sodium L-lactate treatment attenuates the pro-inflammatory activation state of microglia, as reflected by reduced CD86 expression and TNF-α levels, while also decreasing the production of chemokines involved in the recruitment of peripheral immune cells. These findings are consistent with earlier work showing that L-lactate (10-30 mM) suppresses LPS-induced inflammatory responses in microglia (Liang L, *et al*., 2024). Regarding CD86, our results are consistent with an *in vivo* study on mouse tissues (Kong L, *et al*., 2019). Chemokines, including CINC-1/CXCL1 and TIMP-1, are consistently reported to be upregulated in *in vivo* models of cerebral hypoxia-ischemia, both in adults and neonates. Multiple studies have demonstrated that the expression of specific chemokines is significantly increased in the brain following hypoxic-ischemic injury or LPS stimulation. This upregulation underscores their pivotal role in neutrophil recruitment and the amplification of neuroinflammatory responses. The secretion of these chemokines is closely correlated with the extent of brain damage and the polarization of local immune cells, such as macrophages and microglia, toward a pro-inflammatory phenotype. Consequently, these molecules serve as potential biomarkers of inflammation severity and represent promising therapeutic targets in neonatal encephalopathies (Ziemka-Nalecz M, *et al*., 2017). In this context, CINC-1 has been clearly identified as a key biomarker of the neuroinflammatory response following NHI and/or LPS exposure, with robust evidence in neonatal models (Brochu ME, *et al*., 2011). Further supporting the involvement of chemokines in this pathological process, Bednarek *et al*. demonstrated that following hypoxia-ischemia, levels of MMP-9 (matrix metalloproteinase-9) and TIMP-1 significantly increase in both the brain and plasma (Bednarek N, *et al*., 2012). These molecules play critical roles in extracellular matrix degradation, blood-brain barrier (BBB) permeability, and the post-ischemic inflammatory response, further linking chemokine activity to the progression of neuroinflammatory damage. Their increased expression contributes to the establishment of a chemotactic gradient, recruiting additional immune cells to the site of injury and thus participating in the post-traumatic inflammatory responses. Our study demonstrates that *in vitro* sodium L-lactate significantly reduces the expression of key chemokines, including CINC-1, CINC-2, CINC-3, TIMP-1, and MIP-1α. This result suggests that lactate may not only modulate intracellular inflammatory signaling pathways but may also attenuate the microglia-mediated recruitment of peripheral immune cells *in vivo*. Hong *et al*. further demonstrated, using primary microglia, *in vivo* mouse models, and Akt inhibition, that lactate promotes microglial process elongation, prevents LPS-induced retraction, and reduces both inflammatory signaling and associated behavioral disturbances; all these effects required Akt activity (Hong H, *et al*., 2023). Likewise, lactate inhibited NF-κB activation and cytokine release in primary mouse macrophages, human monocytes, and *in vivo* models of acute liver injury and pancreatitis, through a G-protein coupled receptor 81 (GPR81)-dependent mechanism (Hoque R, *et al*., 2014). Together, these *in vitro* and *in vivo* studies corroborate the anti-inflammatory profile observed in our primary microglia experiments.

The mechanisms underlying this effect may involve the close relationship between microglial metabolism and inflammatory activation. Lactate engages key metabolic regulators that couple glycolysis to inflammatory gene expression. Several studies have identified the MCT1/PFKFB3/HIF-1α (monocarboxylate transporter 1/6-phosphofructo-2-kinase/fructose-2,6-bisphosphatase 3/hypoxia-inducible factor 1 alpha) axis and glycolysis-dependent NF-κB activation as central drivers of classical microglial activation. *In vitro* and *in vivo* work has demonstrated that MCT1 enhances glycolytic flux by promoting PFKFB3 through HIF-1α signaling; silencing MCT1 decreases glycolysis and reduces LPS-induced inducible nitric oxide synthase (iNOS), interleukin (IL)-1β, IL-6, and signal transducer and activator of transcription 1 (STAT1) phosphorylation in BV2 microglia (Kong L, *et al*., 2019). Lactate treatment inhibited MCT1-mediated glycolysis, reduced PFKFB3 expression, suppressed pro-inflammatory microglial polarization *in vitro*, and ameliorated LPS-induced sickness behavior *in vivo*, highlighting lactate as a potential anti-inflammatory therapy. Thus, lactate shapes microglial phenotype through its impact on transporter activity, metabolic flux, and downstream signaling pathways.

Pro-inflammatory activation profoundly reshapes microglial metabolism. In BV-2 cells, LPS exposure elevates lactate output while reducing mitochondrial respiration and ATP synthesis, thereby favoring a glycolytic profile that supports enhanced transcriptional activity (Voloboueva LA, *et al*., 2013). Consistent with this shift, primary microglia show altered inflammatory responses (reduced TNF-α and IL-6 production) when glycolysis is blocked with 2-deoxy-D-glucose, though this metabolic inhibition compromises cell viability (Vilalta A and Brown GC, 2014; Wang Q, *et al*., 2014). Moreover, an oversupply of glucose has been shown to amplify TNF-α release in primary rat microglia (Quan Y, *et al*., 2011), further supporting a link between glucose availability, glycolytic activity, and inflammatory activation. Several inflammatory stimuli, including LPS, amyloid-β, and interferon gamma (IFNγ), increase the expression of key glycolytic enzymes such as PFKFB3, hexokinase II, and pyruvate kinase M2 (PKM2), together with elevations in extracellular acidification rate, demonstrating that inflammation drives a glycolytic bias (Rubio-Araiz A, *et al*., 2018). Upregulation of GLUT1 under inflammatory conditions further sustains glycolytic flux, and blocking this transporter can redirect microglia toward oxidative metabolism, dampen activation, and mitigate neurodegeneration *in vivo* (Wang L, *et al*., 2019). Overall, microglial inflammatory activation is closely tied to a shift toward glycolysis, which adapts energy production to the heightened metabolic demands of reactive cells. Rather than simply reflecting a consequence of inflammation, this metabolic reprogramming can contribute to the maintenance of the pro-inflammatory state. Since glycolytic activation is a key driver of pro-inflammatory microglial responses, any intervention capable of counteracting this metabolic shift would be expected to attenuate inflammatory markers. In line with this well-established metabolic reprogramming, LPS stimulation in our primary microglial cultures induced a robust pro-inflammatory phenotype, reflected by the upregulation of CD86 and marked cytokines and chemokines release. Importantly, post-LPS administration of sodium L-lactate attenuated this inflammatory response and reduced these markers, consistent with lactate’s capacity to counteract glycolytic activation described in previous studies. These findings support the concept that lactate can modulate microglial metabolism to limit glycolysis-associated pro-inflammatory activation.

This anti-inflammatory effect of exogenous sodium L-lactate is further supported by studies showing that metabolic interventions that reduce excessive glycolytic reliance can attenuate microglial inflammatory responses. Inducing a ketogenic-like state in microglia, limiting glycolytic flux, has been shown to reduce inflammatory signaling, mitigate tissue damage, and improve functional outcomes after central nervous system injury (Fu SP, *et al*., 2015; Huang C, *et al*., 2018). A key mechanism involves activation of the G-protein–coupled receptor 109A (GPR109A) by the ketone body β-hydroxybutyrate, which suppresses NF-κB-dependent cytokine production and promotes a neuroprotective microglial phenotype *in vivo* (Fu SP, *et al*., 2015; Huang C, *et al*., 2018; Rahman M, *et al*., 2014). These findings are particularly relevant in the context of our results because ketone bodies and lactate share monocarboxylate transporters and function as alternative cerebral energy substrates. Thus, converging evidence indicates that metabolic substrates can directly influence microglial inflammatory states, aligning well with the anti-inflammatory effects we observed with post-LPS lactate treatment.

*In vivo*, sodium L-lactate administered post-insult significantly limited microglial activation 48 h after NHI, as evidenced by normalization of Iba1 levels and microglial density in the cortex and by a partial reduction of these parameters in the hippocampus. Notably, lactate fully restored cortical CD206 expression and prevented the upregulation of CD86, highlighting a shift toward an anti-inflammatory phenotype. In contrast, hippocampal microglia displayed only partial responsiveness, with persistent alterations in both CD86 and CD206 expression. In line with these molecular and phenotypic findings, the analysis of microglial morphology further revealed region-specific effects of lactate. In the cortex, NHI induced a pronounced reduction in microglial arborization complexity, reflected by a decrease in the number of branches, junctions, and branching points. Sodium L-lactate post-treatment fully prevented these structural alterations, preserving a highly ramified morphology consistent with a non-activated or pro-resolutive state. Conversely, hippocampal microglia exhibited a marked loss of complexity following NHI, with reductions across all morphological parameters. Importantly, lactate failed to restore this complexity, indicating that hippocampal microglia remain structurally and functionally impaired despite treatment.

This anti-inflammatory role of lactate observed in the cortex is consistent with several previous studies. Beyond its direct effects on cellular metabolism, lactate may also exert its immunomodulatory actions through several signaling mechanisms. One potential pathway involves the lactate receptor HCAR1 (also known as GPR81) (Kennedy L, *et al*., 2022). Importantly, this study did not involve exogenous lactate administration but rather investigated the role of endogenous lactate signaling following NHI. HCAR1 knockout mice displayed reduced tissue regeneration compared with wild-type animals following NHI. Moreover, neural progenitor and glial proliferation, as well as microglial activation, were altered in the absence of this receptor. These findings suggest that lactate produced endogenously in response to hypoxia–ischemia may itself contribute to the brain’s endogenous neuroprotective response through HCAR1 signaling. Thus, the beneficial effects of lactate may not only result from exogenous sodium L-lactate administration, but may also reflect and reinforce an endogenous lactate-dependent protective mechanism activated following NHI. Lactate may also modulate inflammation through histone lactylation (Zhou Y, *et al*., 2022). Finally, beyond the neonatal HI context, additional evidence from ischemia models further supports a direct immunomodulatory action of lactate on microglia. For example, in a mouse model of cerebral ischemia–reperfusion induced by MCAO, lactate was delivered intracerebroventricularly and microglial responses were quantified 24 h later using immunomarkers, cytokine profiling, and NF-κB signaling assays. The treatment reduced infarct size and neuronal apoptosis while shifting microglia toward an anti-inflammatory profile, decreasing pro-inflammatory markers and increasing anti-inflammatory markers. Mechanistically, lactate acted through HIF-1α, which inhibited NF-κB, and blocking either pathway abolished its anti-inflammatory and neuroprotective effects (Zhang Y, *et al*., 2025).

The difference in neuroprotection observed between the cortex and the hippocampus after post-HI sodium L-lactate administration may be partially explained by intrinsic morphological and vascular differences between these two brain regions. Indeed, regional variations in capillary density have been reported in the rat brain (Borowsky IW and Collins RC, 1989). The cortical capillary network is substantially more developed than that of the hippocampus, a structural characteristic that may favor better delivery and utilization of exogenous sodium L-lactate in the cortex. This vascular disparity may also contribute to the well-established heightened vulnerability of the hippocampal CA1 region to ischemic injury (Ianevski A, *et al*., 2025; Kreisman NR, *et al*., 2000; Lana D, *et al*., 2020; Schmidt-Kastner R, 2015). Consistent with this notion, Cavaglia *et al*. demonstrated that cortical and hippocampal areas display robust, region-specific differences in microvascular organization, including lower capillary density in CA1 compared with cortical regions and even neighboring hippocampal subfields such as CA3 (Cavaglia M, *et al*., 2001). Their work further showed that CA1 vessels exhibit greater BBB disruption following ischemic insult, reinforcing the idea that vascular architecture critically shapes regional susceptibility to injury. Together with our findings, these data suggest that the limited responsiveness of hippocampal microglia to lactate may result, at least in part, from intrinsic constraints related to hippocampal vascularization and its well-documented ischemia-prone nature.

In summary, our results demonstrate that lactate acts as an immunomodulatory signal capable of limiting microglial activation after NHI. However, its efficacy is region-dependent, likely reflecting structural, metabolic, and microvascular differences between cortical and hippocampal microglia. Importantly, lactate did not appear to abolish microglial responses but rather attenuated the excessive pro-inflammatory phenotype while preserving or restoring features associated with a less reactive, potentially pro-resolutive state. This distinction is particularly relevant in the developing brain, where microglial activation is required for physiological functions including synaptic remodeling, tissue repair, and recovery. Additionally, lactate’s emerging role in promoting microglial metabolic flexibility suggests further therapeutic potential for brain disorders, highlighting its importance in both healthy and diseased states (Monsorno K, *et al*., 2022). Understanding how these factors shape microglial responsiveness to metabolic interventions may help optimize therapeutic strategies for neonatal brain injury.

## Conclusion

In conclusion, our study demonstrates that lactate acts as a potent immunometabolic modulator, limiting microglial activation after neonatal hypoxic-ischemic injury in a region-dependent manner. While cortical microglia are highly responsive, hippocampal microglia display limited sensitivity, likely due to intrinsic vascular, metabolic, and structural constraints. These findings highlight the therapeutic potential of targeting microglial metabolism to limit excessive neuroinflammation while preserving the physiological functions of microglia in the developing brain.

## Acknowledgements

The present study was conducted in the context of the framework of the University of Bordeaux’s IdEx‘Investments for the Future’ program/RRI“IMPACT.” This work was supported by the Fédération pour la Recherche sur le Cerveau (FRC) and by the Nouvelle Aquitaine region. This work received also financial support from the French Agence Nationale de la Recherche (ANR) grant, BrainFuel reference (grant number ANR-21-CE44-0023 to A.K.B.S. and L.P.) and the Spark program.

## Statements and declarations

Authors have no conflict of interest to declare.

## Author contribution

JLT, HR and AKBS contributed to the study conception and design. Experiments, data collection and analyses were performed by IOI, JLT and PG. The first draft of the manuscript was written by HR and JLT. All authors commented on previous versions of the manuscript. All authors read and approved the final manuscript.

## References

Bednarek N, Svedin P, Garnotel R, Favrais G, Loron G, Schwendiman L, Hagberg H, Morville P, et al. (2012), Increased MMP-9 and TIMP-1 in mouse neonatal brain and plasma and in human neonatal plasma after hypoxia-ischemia: a potential marker of neonatal encephalopathy. Pediatr Res 71:63–70.

Bernier LP, York EM, MacVicar BA (2020), Immunometabolism in the Brain: How Metabolism Shapes Microglial Function. Trends Neurosci 43:854–869.

Borowsky IW, Collins RC (1989), Metabolic anatomy of brain: a comparison of regional capillary density, glucose metabolism, and enzyme activities. J Comp Neurol 288:401–413.

Brochu ME, Girard S, Lavoie K, Sebire G (2011), Developmental regulation of the neuroinflammatory responses to LPS and/or hypoxia-ischemia between preterm and term neonates: An experimental study. J Neuroinflammation 8:55.

Cavaglia M, Dombrowski SM, Drazba J, Vasanji A, Bokesch PM, Janigro D (2001), Regional variation in brain capillary density and vascular response to ischemia. Brain Res 910:81–93.

Dommergues MA, Plaisant F, Verney C, Gressens P (2003), Early microglial activation following neonatal excitotoxic brain damage in mice: a potential target for neuroprotection. Neuroscience 121:619–628.

Ekdahl CT, Kokaia Z, Lindvall O (2009), Brain inflammation and adult neurogenesis: the dual role of microglia. Neuroscience 158:1021–1029.

Faustino JV, Wang X, Johnson CE, Klibanov A, Derugin N, Wendland MF, Vexler ZS (2011), Microglial cells contribute to endogenous brain defenses after acute neonatal focal stroke. J Neurosci 31:12992–13001.

Fleiss B, Van Steenwinckel J, Bokobza C, I KS, Ross-Munro E, Gressens P (2021), Microglia-Mediated Neurodegeneration in Perinatal Brain Injuries. Biomolecules 11.

Fu SP, Wang JF, Xue WJ, Liu HM, Liu BR, Zeng YL, Li SN, Huang BX, et al. (2015), Anti-inflammatory effects of BHBA in both in vivo and in vitro Parkinson’s disease models are mediated by GPR109A-dependent mechanisms. J Neuroinflammation 12:9.

Gimeno-Bayon J, Lopez-Lopez A, Rodriguez MJ, Mahy N (2014), Glucose pathways adaptation supports acquisition of activated microglia phenotype. J Neurosci Res 92:723–731.

Green TRF, Rowe RK (2024), Quantifying microglial morphology: an insight into function. Clin Exp Immunol 216:221–229.

Hagberg H, Gressens P, Mallard C (2012), Inflammation during fetal and neonatal life: implications for neurologic and neuropsychiatric disease in children and adults. Ann Neurol 71:444–457.

Hong H, Su J, Zhang Y, Xu G, Huang C, Bao G, Cui Z (2023), A novel role of lactate: Promotion of Akt-dependent elongation of microglial process. Int Immunopharmacol 119:110136.

Hoque R, Farooq A, Ghani A, Gorelick F, Mehal WZ (2014), Lactate reduces liver and pancreatic injury in Toll-like receptor- and inflammasome-mediated inflammation via GPR81-mediated suppression of innate immunity. Gastroenterology 146:1763–1774.

Huang C, Wang P, Xu X, Zhang Y, Gong Y, Hu W, Gao M, Wu Y, et al. (2018), The ketone body metabolite beta-hydroxybutyrate induces an antidepression-associated ramification of microglia via HDACs inhibition-triggered Akt-small RhoGTPase activation. Glia 66:256–278.

Ianevski A, Camara-Quilez M, Wang W, Suganthan R, Hildrestrand G, Grini JV, Doskeland DS, Ye J, Bjoras M (2025), Early transcriptional responses reveal cell type-specific vulnerability and neuroprotective mechanisms in the neonatal ischemic hippocampus. Acta Neuropathol Commun 13:147.

Jiao M, Li X, Chen L, Wang X, Yuan B, Liu T, Dong Q, Mei H, Yin H (2020), Neuroprotective effect of astrocyte-derived IL-33 in neonatal hypoxic-ischemic brain injury. J Neuroinflammation 17:251.

Jin D, Dai Z, Zhao L, Ma T, Ma Y, Zhang Z (2024), CYR61 is Involved in Neonatal Hypoxic-ischemic Brain Damage Via Modulating Astrocyte-mediated Neuroinflammation. Neuroscience 552:54–64.

Jurga AM, Paleczna M, Kuter KZ (2020), Overview of General and Discriminating Markers of Differential Microglia Phenotypes. Front Cell Neurosci 14:198.

Kennedy L, Glesaaen ER, Palibrk V, Pannone M, Wang W, Al-Jabri A, Suganthan R, Meyer N, et al. (2022), Lactate receptor HCAR1 regulates neurogenesis and microglia activation after neonatal hypoxia-ischemia. Elife 11.

Kong L, Wang Z, Liang X, Wang Y, Gao L, Ma C (2019), Monocarboxylate transporter 1 promotes classical microglial activation and pro-inflammatory effect via 6-phosphofructo-2-kinase/fructose-2, 6-biphosphatase 3. J Neuroinflammation 16:240.

Kreisman NR, Soliman S, Gozal D (2000), Regional differences in hypoxic depolarization and swelling in hippocampal slices. J Neurophysiol 83:1031–1038.

Lalancette-Hebert M, Gowing G, Simard A, Weng YC, Kriz J (2007), Selective ablation of proliferating microglial cells exacerbates ischemic injury in the brain. J Neurosci 27:2596–2605.

Lana D, Ugolini F, Giovannini MG (2020), An Overview on the Differential Interplay Among Neurons-Astrocytes-Microglia in CA1 and CA3 Hippocampus in Hypoxia/Ischemia. Front Cell Neurosci 14:585833.

Li B, Concepcion K, Meng X, Zhang L (2017), Brain-immune interactions in perinatal hypoxic-ischemic brain injury. Prog Neurobiol 159:50–68.

Liang L, Liu P, Deng Y, Li J, Zhao S (2024), L-lactate inhibits lipopolysaccharide-induced inflammation of microglia in the hippocampus. Int J Neurosci 134:45–52.

Liddelow SA, Barres BA (2017), Reactive Astrocytes: Production, Function, and Therapeutic Potential. Immunity 46:957–967.

Liddelow SA, Olsen ML, Sofroniew MV (2024), Reactive Astrocytes and Emerging Roles in Central Nervous System (CNS) Disorders. Cold Spring Harb Perspect Biol 16.

Liddelow SA, Sofroniew MV (2019), Astrocytes usurp neurons as a disease focus. Nat Neurosci 22:512–513.

Lin L, Desai R, Wang X, Lo EH, Xing C (2017), Characteristics of primary rat microglia isolated from mixed cultures using two different methods. J Neuroinflammation 14:101.

Milicevic K, Korenic A, Milosevic M, Andjus PR (2022), Primary Cultures of Rat Astrocytes and Microglia and Their Use in the Study of Amyotrophic Lateral Sclerosis. J Vis Exp.

Millar LJ, Shi L, Hoerder-Suabedissen A, Molnar Z (2017), Neonatal Hypoxia Ischaemia: Mechanisms, Models, and Therapeutic Challenges. Front Cell Neurosci 11:78.

Monsorno K, Buckinx A, Paolicelli RC (2022), Microglial metabolic flexibility: emerging roles for lactate. Trends Endocrinol Metab 33:186–195.

Nimmerjahn A, Kirchhoff F, Helmchen F (2005), Resting microglial cells are highly dynamic surveillants of brain parenchyma in vivo. Science 308:1314–1318.

Quan Y, Jiang CT, Xue B, Zhu SG, Wang X (2011), High glucose stimulates TNFalpha and MCP-1 expression in rat microglia via ROS and NF-kappaB pathways. Acta Pharmacol Sin 32:188–193.

Rahman M, Muhammad S, Khan MA, Chen H, Ridder DA, Muller-Fielitz H, Pokorna B, Vollbrandt T, et al. (2014), The beta-hydroxybutyrate receptor HCA2 activates a neuroprotective subset of macrophages. Nat Commun 5:3944.

Roumes H, Dumont U, Sanchez S, Mazuel L, Blanc J, Raffard G, Chateil JF, Pellerin L, Bouzier-Sore AK (2021), Neuroprotective role of lactate in rat neonatal hypoxia-ischemia. J Cereb Blood Flow Metab 41:342–358.

Roumes H, Pellerin L, Bouzier-Sore AK (2020), Neuroprotective role of lactate in neonatal hypoxia-ischemia. Med Sci (Paris) 36:973–976.

Rubio-Araiz A, Finucane OM, Keogh S, Lynch MA (2018), Anti-TLR2 antibody triggers oxidative phosphorylation in microglia and increases phagocytosis of beta-amyloid. J Neuroinflammation 15:247.

Schmidt-Kastner R (2015), Genomic approach to selective vulnerability of the hippocampus in brain ischemia-hypoxia. Neuroscience 309:259–279.

Sen E, Levison SW (2006), Astrocytes and developmental white matter disorders. Ment Retard Dev Disabil Res Rev 12:97–104.

Serdar M, Kempe K, Rizazad M, Herz J, Bendix I, Felderhoff-Muser U, Sabir H (2019), Early Pro-inflammatory Microglia Activation After Inflammation-Sensitized Hypoxic-Ischemic Brain Injury in Neonatal Rats. Front Cell Neurosci 13:237.

Sullivan SM, Bjorkman ST, Miller SM, Colditz PB, Pow DV (2010), Morphological changes in white matter astrocytes in response to hypoxia/ischemia in the neonatal pig. Brain Res 1319:164–174.

Vilalta A, Brown GC (2014), Deoxyglucose prevents neurodegeneration in culture by eliminating microglia. J Neuroinflammation 11:58.

Voloboueva LA, Emery JF, Sun X, Giffard RG (2013), Inflammatory response of microglial BV-2 cells includes a glycolytic shift and is modulated by mitochondrial glucose-regulated protein 75/mortalin. FEBS Lett 587:756–762.

Wang L, Pavlou S, Du X, Bhuckory M, Xu H, Chen M (2019), Glucose transporter 1 critically controls microglial activation through facilitating glycolysis. Mol Neurodegener 14:2.

Wang Q, Zhao Y, Sun M, Liu S, Li B, Zhang L, Yang L (2014), 2-Deoxy-d-glucose attenuates sevoflurane-induced neuroinflammation through nuclear factor-kappa B pathway in vitro. Toxicol In Vitro 28:1183–1189.

Zhang Y, Zhang S, Yang L, Zhang Y, Cheng Y, Jia P, Lv Y, Wang K, et al. (2025), Lactate modulates microglial inflammatory responses through HIF-1alpha-mediated CCL7 signaling after cerebral ischemia in mice. Int Immunopharmacol 146:113801.

Zhou Y, Yang L, Liu X, Wang H (2022), Lactylation may be a Novel Posttranslational Modification in Inflammation in Neonatal Hypoxic-Ischemic Encephalopathy. Front Pharmacol 13:926802.

Ziemka-Nalecz M, Jaworska J, Zalewska T (2017), Insights Into the Neuroinflammatory Responses After Neonatal Hypoxia-Ischemia. J Neuropathol Exp Neurol 76:644–654.

